# The insect brain is specified by a conserved transcriptional program distinct from the segmental CNS

**DOI:** 10.64898/2026.08.27.747565

**Authors:** Noel Cabañas, Ana Veloso, Robert Zinzen, Gregor Bucher

**Author notes:** **Corresponding author**: Gregor Bucher, Justus-von-Liebig Weg 11, 37077 Göttingen.

## Abstract

The brain is the central processing center that integrates information to guide animal behavior, while the posterior CNS performs less central functions. To what degree this functional distinction is reflected in divergence of the genetic regulation remains unexplored, as well as the hypothesis of a distinct evolutionary origin of brain and trunk CNS. In insects, neuroblast biology is similar across the CNS, but the presence of brain-specific neuroblast types and differences in the gene sets involved in their spatial and temporal specification indicate divergence. However, a comprehensive and unbiased comparison of the regulatory code of brain versus trunk neuroblasts has been lacking. Here, we reveal the transcription factors expressed in insect embryonic neuroblasts by a combined single-cell analysis from the fly *Drosophila melanogaster* and the beetle *Tribolium castaneum,* for which we established a Gal4 enhancer trap system to generate a line marking NBs with tGFP. From the combined dataset of nearly 70,000 neuroblasts, we identified around 23,000 brain neuroblasts, from which we identified cell populations and their transcriptional signatures. We found 188 brain-specific TFs, including seven spatially restricted transcription factors not previously associated with neuroblasts (*Hmx, CG15CSC, CG32532, dmrtSSB, fD5SA, TfAP-2*, and *Fer1*). In addition to the expected canonical Type II NBs, we found two related clusters with intriguing transcriptional divergence. The large number of brain-specific TFs supports a profound divergence between the insect brain and posterior CNS. Together with recent work in vertebrates, our data confirm an ancient divergence of the brain from the trunk.

**Significance statement:** The brain integrates information to guide animal behaviour while the rear parts of the nervous system act mainly downstream. Although function and structures of the brain are quite distinct, they start their development in a similar way. Therefore, it has remained unclear how different the genetic regulation of the brain actually is. To answer this, we revealed the genetic program of brain stem cells at single-cell resolution across two insects — a fly and a beetle. We comprehensively identified key genetic regulators active in the brain but not the posterior nervous system, including several previously unknown ones. This divergence suggests that the brain is quite fundamentally different from the trunk CNS — a finding which resonates with recent studies from vertebrates.

## Introduction

The brain is essential for the survival of an animal as it governs behaviour by integrating external sensory information with internal physiological states and memory. The more posterior parts of the central nervous system (CNS) are more involved in the realization of the decisions taken in the brain. This functional divergence is reflected in brain-specific neural structures and by the expression of transcription factors involved in anterior neural development of all animals (1–4). It was even suggested that anterior and posterior parts of the CNS evolved independently and fused to form a “chimeric brain” (5, 6). On the other side, basic cellular and genetic mechanisms of CNS development are quite similar in both regions, arguing more for a common origin of the CNS with subsequent elaboration and specification of the brain. An unbiased comparison of the divergence of the genetic control of brain and posterior CNS could shed light on this issue but has been lacking in any arthropod.

Insects are excellent model systems for addressing evolutionary questions, as they display a great degree of brain diversification and are amenable to genetic manipulation. The insect CNS consists of the posterior segmental ventral nerve cord (VNC) and the brain, which has both segmental and non-segmental regions (4, 7). It develops from neural stem cells, named neuroblasts (NBs), that delaminate from the ventral and procephalic neuroectoderm into the embryo before they begin proliferating (8, 9). Before delamination, the NBs obtain their spatial identity by the expression of “cocktails” of transcription factors, which are well-described for the VNC but poorly understood for the brain(5, 8–13).

Two main types of NBs are distinguished. Type I NBs that divide several times asymmetrically to produce a self-renewing NB and ganglion mother cells (GMCs) that divide once more to produce two neural cells. The Type I proliferation mode is found in the VNC NBs and in most brain NBs. Type II NBs are found only in the brain, where they divide asymmetrically to produce several intermediate neural progenitors (INPs) that divide in a stem cell-like fashion as well to produce GMCs, which divide once more to produce neural cells. Therefore, Type II NBs produce much larger neural lineages (10, 14–16).

The genetic mechanisms that determine the fate of a neuroectodermal cell to become an NB are well understood and are similar for NBs from both VNC and brain, and are conserved in insects (8, 9, 11, 17, 18). Each NB exhibits a distinct identity and produces a unique and stereotypical neural lineage (19, 20). NBs obtain their individual identity in a stepwise patterning process. The first signal is spatial information that NBs inherit from their location of origin within the neuroectoderm. In the VNC of flies, the spatial information consists of the grid-like expression of segment polarity and columnar genes, along with the regionally restricted Hox gene expression. The different combinations of these transcription factors during NB delamination make up a “cocktail” of transcription factors (TFs) that confers a unique identity on each NB (8, 9). Subsequent diversifying signals are provided by temporal transcription factor cascades, which lead to the specification of different neurons depending on their time of birth (8, 11, 20). Functionally, the spatial transcription factors seem to modulate the open chromatin architecture of NBs, which in turn modifies the NB’s responsiveness to the expression of subsequent temporal transcription factors (21).

In contrast to the VNC, the spatial transcription factor code of brain NBs remains poorly understood. The insect brain is composed of three ganglia, where the posterior two derive from segments, which are serially homologous to trunk segments. The tritocerebrum develops from the intercalary segment and the deutocerebrum from the antennal segment. Consistent with serial homology, the same set of genes is suggested to be involved in their neural development with some variation (22). The pre-antennal part of the CNS is different in several respects. It harbours unique higher order brain structures such as the *mushroom bodies*, the *central complex*, and the ocular lobes, which have no homologs in segmental ganglia. The protocerebrum is built by about 80-100 NBs, roughly three times the number found in the trunk ganglia (13), and it is the only neuromere containing Type II NBs (16, 23–25). At the molecular genetic level, the protocerebrum develops in a region where Hox cluster genes are not expressed and where both, segment polarity and columnar genes, are expressed in divergent patterns or not at all (4, 26). Transcription factors known from VNC patterning have been used as markers for insect brain NBs (12, 13, 27, 28), including only a small subset of the brain-specific aGRN genes. The few aGRN transcription factors that have been functionally studied in insects are essential for protocerebrum-specific structures such as *eyeless* (eyes and *mushroom bodies*), *orthodenticle* (ocular region) (29–31), *six3/Optix* (anterior neuroendocrine part of the brain and *central complex*) (32, 33), *foxQ2* (*mushroom bodies* and *central complex*) (34), and *retinal homeobox* (*mushroom bodies* and *central complex*) (35–38). In summary, the transcription factor code for brain neuroblast specification is at least partially different from the trunk, brain specific TFs contribute to the development of key structures of the brain, yet they remain poorly studied (39), and the temporal cascade in NBs of the optic medulla and the type II NBs diverges from the cascade known from the trunk (8, 40–43). Indeed, molecular and cellular divergence found in animals has been taken as evidence for separate evolutionary origins of brain and posterior CNS (5, 6). However, a comprehensive inventory of TFs differentially expressed in insect brain NBs has been conspicuously absent.

Single-cell RNA sequencing (scRNA-seq) has emerged as a powerful technique for gene expression profiling at cellular resolution, providing a new approach to identify TFs expressed in certain cell types (44, 45). The vinegar fly *Drosophila melanogaster* is the prime candidate for such an endeavour because it is technically the most powerful genetic model system within insects and has been used for pioneering research into CNS development and function (46, 47). Previously, subsets of larval NBs have been studied by scRNA-seq such as the ocular and the Type II NBs (40, 41, 48). However, fly head and neuroectoderm development are extremely derived within insects, (7) and its 106 procephalic NBs are fewer compared to other species such as the beetle *Tenebrio molitor*, which has 124-131 NBs (13, 49). Furthermore, flies have eight Type II NBs per hemisphere while *Tribolium* has nine (16, 23, 24). In contrast to flies, the red flour beetle *Tribolium castaneum* exhibits an insect-typical head development and is an emerging genetic model system with an advanced toolkit, including transgenesis, genome editing for generating neural imaging lines, and systemic RNAi, which has allowed for genome-wide RNAi screens (34, 50–57).

To reveal the degree of divergence of brain versus VNC patterning and to obtain a comprehensive view of the TFs involved in spatial brain NB patterning of insects, we compared single cell transcriptomes of embryonic neuroblasts from two evolutionarily divergent holometabolous insects, *Drosophila* and *Tribolium*. We identified a repertoire of transcription factors differentially expressed in insect brain NBs compared to VNC NBs and validated our dataset by describing seven TFs previously not associated with brain NBs. The large degree of divergence in the TFs active in brain and VNC NBs argues for a profoundly different patterning based on an ancient evolutionary separation. Beyond the evolutionary insight, the transcriptional program revealed by our dataset provides the foundation for understanding the ancestral developmental program on top of which molecular adaptations are deployed and combined to generate the brain diversity across insects.

## Results

### Single-cell neuroblast atlas of two model insects: *Drosophila* and *Tribolium*

#### Isolation of fly neuroblast nuclei and beetle neuroblasts for sequencing

In insects, all cells of the CNS stem from NBs **(Figure 1A)**, which have unique molecular identities that determine the development of their lineages. However, the “cocktails” of TFs responsible for conferring identity to brain NBs remain largely unknown. To gain this information, we performed single cell sequencing in the vinegar fly, *Drosophila melanogaster*, as the pioneering model system of CNS development, and the red flour beetle, *Tribolium castaneum,* as the runner-up with respect to gene function tools. In flies, we used an antibody targeting the *Drosophila* NB marker Worniou to fluorescently mark nuclei of NBs. Nuclei of stained embryos from stages 10 to 12 were collected and enriched using FACS (**Figure supplement 1**). To mark subsets of cells *in vivo* in *Tribolium*, we established a transgenic line with a Gal4 enhancer trap construct. With this, we performed the first Gal4 enhancer trap screen in *Tribolium* by essentially following the procedure of the GEKU enhancer trap screen (see methods and **Figure supplement 2)** (55). We identified a line (GöGal41519), which drove a UAS*-turboGFP* responder in a pattern reminiscent of NBs. Co-expression with the NB marker *asense* confirmed an extensive overlap (**Figure supplement 3**). We then determined the embryonic stages NS9 to NS13 (18) to contain the highest numbers of NBs by inspecting FISH and qPCR of the NB marker *Tc-asense* (**Figure supplement 4A**). We established a procedure to generate cell suspensions from embryonic stages NS9 to NS13 and to enrich for live NBs using FACS (**Figure supplement 4B and C**). We then produced single cell transcriptomic data for both insects using the 10x Genomics technology **(Figure 1B** and **Figure supplement 1 and 4**).

**Figure 1.**
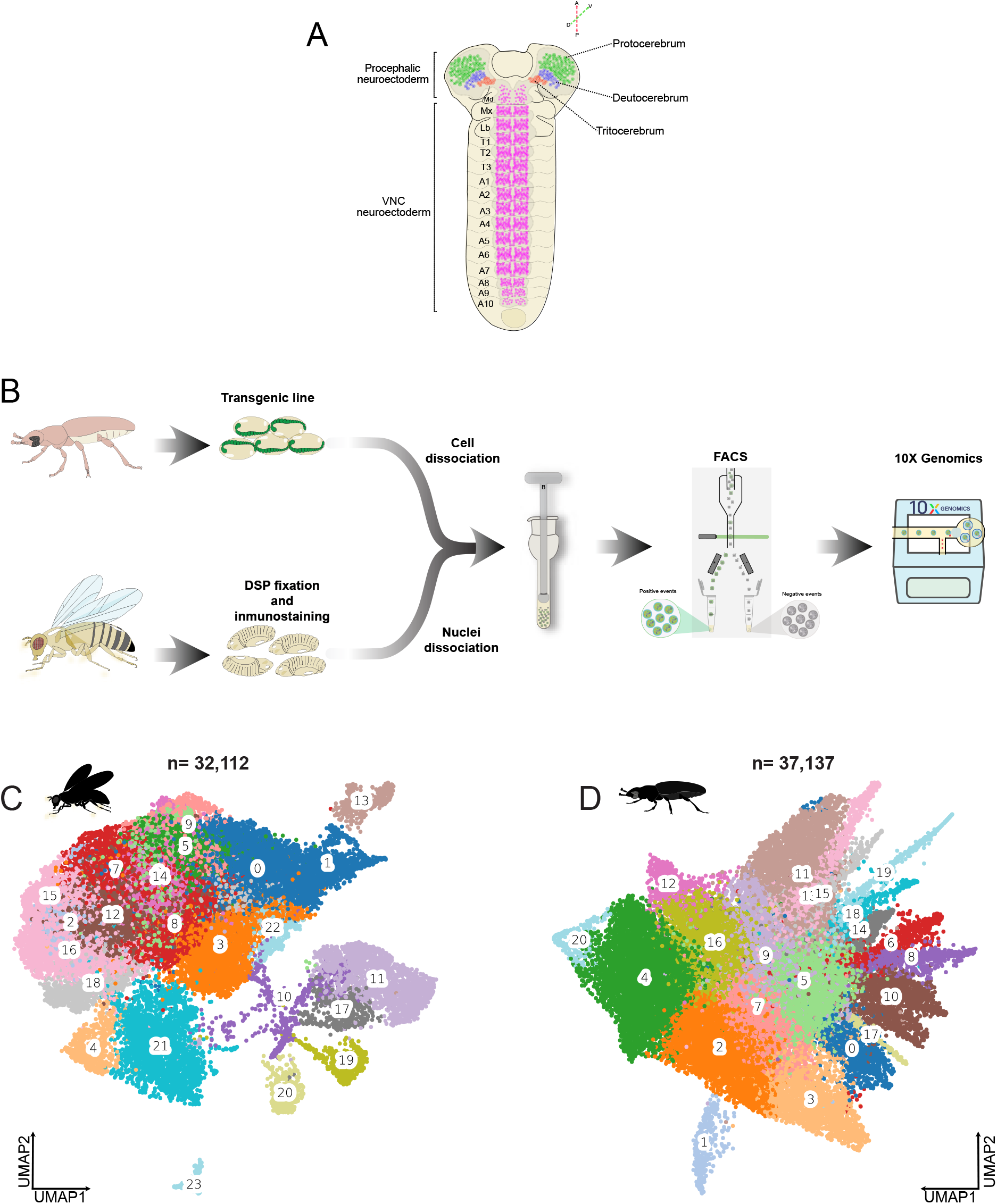
Single-cell atlases of *Tribolium* and *Drosophila* neuroblasts. **(A)** Depicts a representation of a late *Drosophila melanogaster* embryo, with the NBs of the VNC in pink and those of the brain in red, blue and green indicating tritocerebral, deutocerebral, and protocerebral NBs, respectively. **(B)** In *Tribolium*, the workflow started with the isolation of transgenically marked NBs, which were sorted and sequenced using 10x Genomics technology. *Drosophila* NBs were stained with an antibody against Worniou, and nuclei were isolated and sequenced using 10x Genomics technology. **(C)** The UMAP plot of the *Drosophila* atlas with 32,112 nuclei in 24 clusters. **(D)** The *Tribolium* atlas consists of 37,137 cells in 21 clusters. The same pipeline was used for both species in all analyses. Identical colours in these plots do not indicate similarity between fly and beetle.

#### Data processing and quality control

The resulting datasets from both species were processed using the same pipeline to enhance comparability. We started by inspecting the data quality and determined the thresholds for filtering out low-quality reads and empty droplets **(Figure supplements 5-8).** We clustered the cells using an unsupervised approach **(Figure supplements 9-12),** confirmed the expression of the marker genes (*worniou* in *Drosophila* and *tGFP* in *Tribolium*, respectively), assessed the connectivity of the unsupervised clustering, and identified the gene markers in each cluster **(Figure supplements 13-16)**. Further, we validated the presence of marker genes of neural precursors such as *ase*, *wor*, *dpn*, *mira*, *klu*, *erm*, *pnt*, and *pros* **(Figure supplements 17-18)**. As expected, we found these markers to be expressed, but the unsupervised clustering did not reveal clear clusters for subpopulations such as NBs or GMCs. As a negative control, we used the glia marker *repo,* which was not found in the datasets **(Figure supplements 17-18)**. Our final UMAP plots display cell atlases of 32,112 *Drosophila* NB nuclei and 37,137 *Tribolium* embryonic NBs **(Figure 1C, 1D and Figure supplements 9-16)**. These single cell atlases represent a comprehensive transcriptomic resource that contains the molecular profiles of embryonic VNC and brain neuroblasts of both *Drosophila* and *Tribolium*. These species are the most-developed genetic insect model systems and represent two of the four hyperdiverse holometabolan clades. Importantly, these atlases exhibit a high degree of cross-species comparability, owing to the processing with the same pipeline and based on our previous efforts towards robust orthology assignment between these insects. However, for technical and historical reasons, we used different modes of sequencing (nuclei versus live cells), thereby introducing some technical bias.

#### Separating neuroblasts of the ventral nerve cord from the brain

Our atlases present a comprehensive and unbiased overview of the transcriptomic landscape of insect embryonic NBs. NBs derived from serially homologous segments (“VNC NBs”) are expected to share similar TF cocktails. While the specification of segmental NBs has been well studied, the TF cocktails of brain NBs have remained poorly described. Due to our focus on brain NBs, we first sought to differentiate between VNC and brain NBs. To our knowledge, there is no single marker gene distinguishing these cells, but expression of Hox cluster genes (*abd-A*, *Abd-B*, *Antp*, *Scr*, *Dfd*, *pb*, *lab*) is absent from antennal and pre-antennal regions (**Figure 2A, B**). Hence, absence of any Hox gene expression was used to define anterior brain NBs, i.e. those that contribute to the deuto- and protocerebrum. For simplicity, we call these non-Hox NBs “brain NBs”. Note that the tritocerebrum is also considered a part of the brain but was excluded here because it does express the Hox-gene *labial*. The expression of at least one Hox gene was used to classify VNC NBs (**Figure 2C, 2D**). In the *Drosophila* dataset, 10,425 cells were classified as brain NBs and 21,687 as VNC NBs, while in the *Tribolium* dataset, 12,389 cells were brain NBs and 24,748 were VNC NBs **(Figure supplements 17-18)**. Not surprisingly, the distinction between brain versus VNC NBs was not reflected in the clusters found in the entire dataset, because the cell biology of NBs is very similar regardless of the region from which they derive, implying that many genetic markers are expected to be shared across all NBs **(Figure 2E, 2F).**

**Figure 2.**
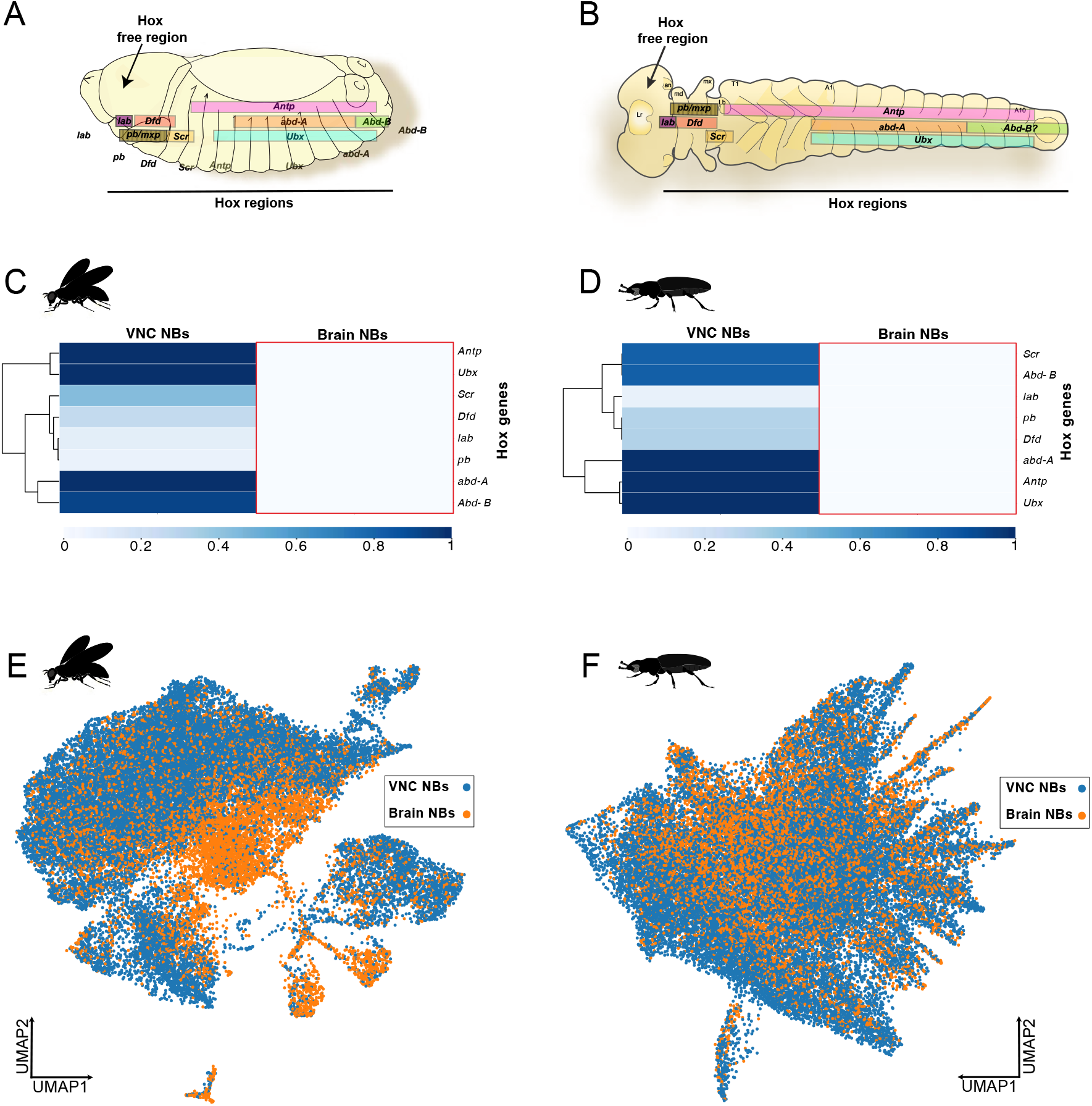
Separation of VNC and brain neuroblasts by virtue of Hox gene expression. (A, **B)** The anterior neuroectoderm in animals is free of the expression of the Hox cluster genes. In insects, the Hox-free neuroectoderm gives rise to proto- and deutocerebrum. **(C, D)** Cells expressing at least one Hox gene were defined as VNC NBs and those lacking any Hox gene expression were defined as brain NBs. **(E, F)** UMAP plots showing that NBs from the brain and VNC do not clearly separate into clusters. Instead, many cells seem to overlap when classifying them as VNC and brain NBs.

#### Manual annotation of brain NBs in *Drosophila* reveals “atypical” Type II NB-like clusters

The protocerebrum is different from the segmental ganglia because it stems from a more evolutionary ancient pre-segmental neuroectoderm and it forms brain-specific higher order processing centres such as the optic lobes, mushroom bodies, and central complex. It is marked by a set of transcription factors with nearly exclusive anterior expression, a feature shared by the anterior brain regions of bilaterians (4). The antennal segment presents an intermediate pattern, where the NBs show similarity to both the trunk segments, but also have molecular and structural divergences that are antenna-specific (58). Specifically, it is the only segment devoid of the Hox gene expression that forms another brain-specific structure, the antennal lobes. To focus on the transcription factors expressed in proto- and deutocerebrum NBs, we subsampled the brain NBs and performed unsupervised clustering. We used the classical gene markers to annotate the *Drosophila* NBs into NBs Type I and GMCs (red and blue in **Figure supplements 19-20**). We also identified a cluster that contained a mix of cells with the expected expression patterns of NBs II and INPs (purple in **Figure supplement 21**). Interestingly, we also identified a couple of atypical clusters that expressed gene markers of both NBs Type II and INPs in addition to some unusual gene markers, so we named them NB II/INPs 1 and 2 (pink and brown clusters in **Figure supplements 21-22**). We also found small clusters of midline NBs and sensory neuroblasts (green and turquoise in **Figure supplements 23 and 24**). Finally, we observed clear clusters of mesodermal/muscle progenitors, PGCs (primordial germ cells), and embryonic hemocytes (**Figure supplements 25-27**).

### Integration of *Tribolium* and *Drosophila* datasets and label transfer

*Drosophila* manual classification was based on the well-described NB marker genes. However, due to some unclear orthology assignments, this approach did not have the same resolution in the *Tribolium* dataset. Therefore, we used the *Drosophila* classification to annotate the *Tribolium* brain NBs. First, we integrated *Tribolium* and *Drosophila* datasets based on orthologous genes using the tool Harmony (59) (**Figure 3A**) and then used scvi-tools and Scanpy to perform the label transfer method from the fly cell clusters to the beetle cells (60, 61). With this approach, we were able to annotate the same cell types in *Tribolium* (**Figure 3C, D**), except for the PGCs. As expected, both datasets were enriched for NBs Type I and GMCs (**Figure 3E**). However, the *Drosophila* dataset had more NB Type I than GMC while in *Tribolium* this was the other way around. This may be due to a technical bias when comparing single-cell and single-nuclei sequencing, or to the differences in marker genes used to enrich the NB cell or nuclei suspensions.

**Figure 3.**
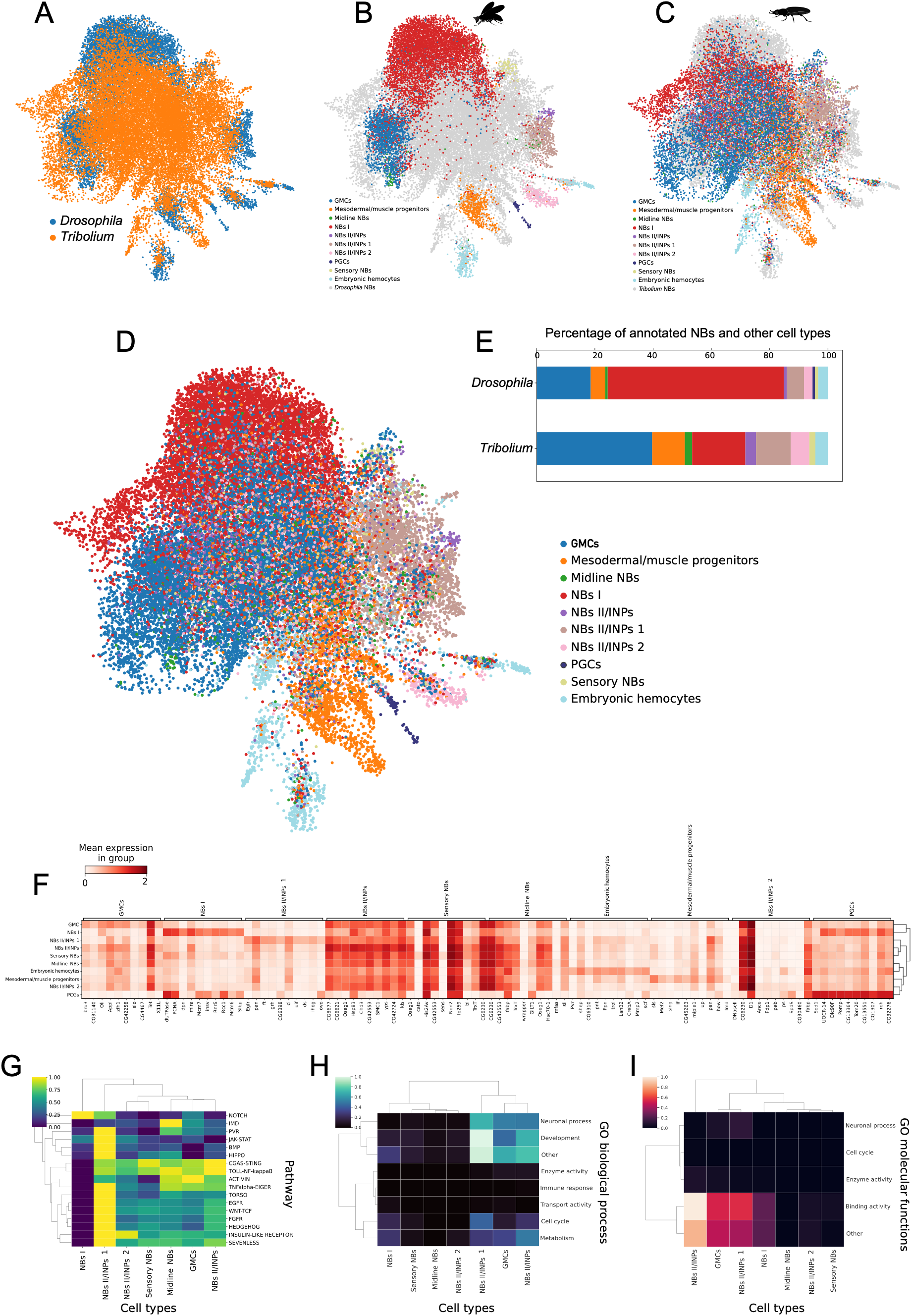
Cell type annotation of *Drosophila* and *Tribolium* brain NBs. **(A)** Plot based on the integrated single cell datasets from *Drosophila* (blue) and *Tribolium* (orange). **(B)** Manual annotation of *Drosophila* NBs using known NB marker genes. *Tribolium* cells are in grey. **(C)** Label transfer of *Drosophila* manual annotations to *Tribolium* NBs. *Drosophila* cells are in grey. Colours in B and C mark the same cell types. **(D)** Integrated dataset that includes both insects and all identified cell types. **(E)** Proportions of cell types revealed a larger number of Type I NBs in the fly data and more GMCs in the beetle data. We assume that this difference is not biological but due to the bias introduced by different isolation techniques and the gene markers used for identifying the NBs. **(F)** The 10 top marker genes for each cell type identified in the integrated dataset. Some clusters are clearly distinguished by these markers (e.g. Type I NBs, PGCs) others are less clearly separated (midline NBs, sensory NBs). Type II NBs marker genes also show expression in most other cell types but are most profoundly different from PGCs. In contrast, Type I NBs show much more similarity with PGCs. **(G)** Pathway enrichment analysis in the NB subtypes differentiates atypical NBs II/INPs 1 from all other clusters by the many involved pathways. Surprisingly, the IMD and cGas-STING immune pathways mark midline NBs and Type II NBs, respectively. **(H)** Gene ontology enrichment analysis for biological processes in broad categories. **(I)** Gene ontology analysis for molecular functions in broad categories.

#### The “atypical” NB Type II-like clusters diverge in signalling pathway and marker gene expression

In the combined dataset, the cluster with the regular NBs II/INPs showed expression of many canonical marker genes at high levels (**Figure 3F**), contrasting the patterns of Type I NBs and the two atypical NBs II/INPs clusters. To further characterize the NB clusters, we performed a signalling pathway enrichment analysis. The Notch pathway is known to remain active after delamination in Type I NBs (62). In Type II NBs, Notch signalling is active as well, but it is downregulated in immature INPs before being re-activated in mature INPs (63). As expected, the Notch pathway was highly active in the Type I NB cluster (**Figure 3G**). However, only the atypical NBs II/INPs 1 showed high expression of Notch components in addition to a surprisingly high number of other signalling pathways (second column from the left in **Figure 3G**). At the same time, it showed a comparably low expression of marker genes (**Figure 3F**), which might indicate that these cells are still quite undifferentiated. The atypical NBs II/INPs 2, in contrast, showed low amounts of Notch pathway components but very high expression of the marker genes CG6230 and D1 (ATP-dependent transporter CG6230 and chromatin binding protein D1) (**Figure 3F**). Hence, the three NBs II/INPs clusters do show puzzling different characteristics, which remain to be understood. Gene ontology analysis revealed that the GMC, NBs II/INPs, and the atypical NBs II/INPs 1 show a clear signal for neuronal processes, development, cell cycle, and metabolism. Only these clusters showed strong enrichment of proteins with specific molecular functions, while the other clusters including atypical NBs II/INPs cluster 2 showed not much expression of genes with GO annotations (**Figure 3H, I**). Surprisingly, we found strong expression of the IMD immune pathway in midline NBs. The expression of PVR pathway components in midline NBs is novel as well, but relates well to its known role in midline glia development (64, 65). We also found that GMCs were more similar in signalling pathway expression to the regular NBs II/INPs than to the atypical Type II clusters or the Type I NBs (**Figure 3G**).

#### Transcription factors differentially expressed in brain and VNC neuroblasts

We wanted to reveal the transcription factors that contribute to the specification of insect brain NB identity using our combined data from fly and beetle. The transcription factor code responsible for VNC NBs is well-understood but respective signals for brain NBs must be different because segment polarity and columnar genes, as well as Hox genes, are expressed either in strongly divergent patterns or not at all in the anterior neuroectoderm, and the number of protocerebral NBs is severalfold higher than in a trunk segment. Some highly conserved transcription factors are expressed in the anterior region of bilaterian animals (4). However, this set represents the most conserved part of the patterning system, which is biased towards those genes that are conserved across Bilateria. An insect-specific set of brain-specific TFs expressed in NBs had not been determined in an unbiased way. To provide a reliable basis for cross-species comparisons, we first predicted orthology of all genes between *Drosophila* and *Tribolium* by using both Orthofinder and EggNogg and included manual quality checks (66–69). Differential expression analysis of all genes based on genome-wide orthology prediction (66) revealed 520 DE genes: 166 were highly enriched in the brain and 354 in the VNC (**Figure supplement 31A**). In *Tribolium*, we found 271 genes: 84 enriched in the brain and 187 in the VNC NBs (**Figure supplement 31B**). As we were interested in the regulatory control of NBs identities, we then focused our analyses on transcription factors. For *Drosophila*, curated lists of TFs exist such as the Jaspar database (141 TFs) and Redfly (304 TFs) (70, 71). More comprehensive lists are available from Flybase (628 TFs) and FlyTF (1052 TFs) (72, 73) and studies from *Drosophila* NBs (15, 41) (**Figure supplement 28**). However, no such effort had been done in *Tribolium* and a simple search for *Tribolium* orthologs of fly TFs would introduce a species bias by missing beetle-specific TFs and by their diverging annotation quality. Hence, to ensure comparability between species, we did a *de novo* prediction of all TFs for *Tribolium* and *Drosophila* using DeepTFactor (74). This resulted in 868 *Tribolium* and 958 *Drosophila* TFs (**Figure supplement 29A, B,** and **C**), which is in the range of the previous estimations in *Drosophila*. We then asked which TFs were differentially expressed in brain versus VNC NBs. In *Drosophila,* 290 TFs were significantly differentially expressed: 153 in the VNC NBs and 137 in the brain NBs (**Figure 4A and Figure supplements 29D - F and G - I**). For *Tribolium*, we found 167 TFs to be differentially expressed: 89 in the VNC and 78 in the brain NBs (**Figure 4B and Figure supplement 29 D, E and G, H).**

**Figure 4.**
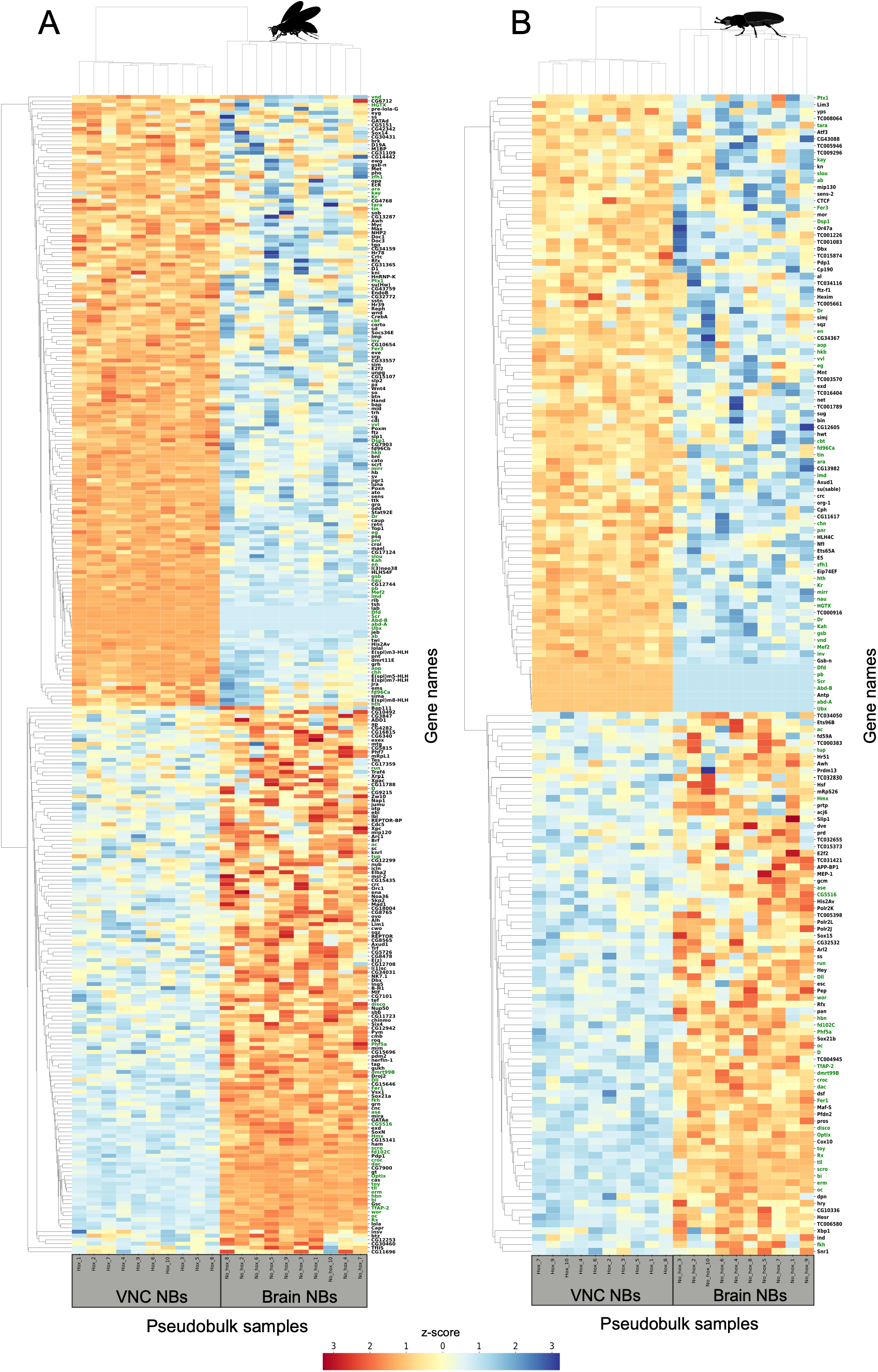
Transcription factors differentially expressed in *Tribolium* and *Drosophila* NBs. We performed independent pseudobulk differential expression analysis to identify significant TFs for VNC and brain NBs in each species. The heatmap plots show 290 and 167 TFs identified in *Drosophila* and *Tribolium* respectively. (**A**) In *Drosophila,* we identified 153 TFs DE in the VNC and 137 in the brain. (**B**) In *Tribolium*, we identified 89 TFs in the VNC and 78 in the brain NBs. Gene names in green represent 27 TFs shared by *Drosophila* and *Tribolium*.

#### Transcription factors enriched in the insect VNC neuroblasts

While our focus was on brain NBs, we also checked the VNC-specific genes. 37 orthologous genes were conserved in their enrichment of expression in VNC NBs of both *Drosophila* and *Tribolium*: *Abd-B, Dfd, Dr, Dsp1, Fer3, HGTX, Kah, Kr, Mef2, Ptx1, Scr, Ubx, ab, abd-A, aop, ara, cbt, chn, eg, en, fd96Ca, gsb, hkb, hth, inv, kay, lmd, mirr, nau, pb, pnr, slou, tara, tin, vnd, vvl, zT1.* Confirming our approach, the list comprises the Hox genes and long-known markers such as *vnd*, *en*, and *hkb* in addition to more recently identified markers such as *fd96Ca* or *Fer3* (75) (**Figure 4**) **(Figure supplement 29)**. Given that many expected genes were identified, the novel genes are very good candidates for future research. However, they first need to be confirmed by whole mount *in situ* hybridization. For instance, we found *Mef2* and *slou* expression in mesodermal cells but not in NBs **(Figure supplements 84-85)** and some genes do show expression in the brain as well, e.g. *en, ems*, *Ptx1* and *slp1*. We assume that the segmental repetition of the NB pattern leads to a much higher number of trunk NBs expressing a certain TF compared to the brain potentially leading to apparent differential expression.

#### Novel transcription factors in insect brain neuroblasts

The brain harbours higher order cognitive abilities such as orientation in space (*central complex*) and learning and memory (*mushroom bodies*). To determine genes potentially involved in the regulatory code of such brain-specific structures, we determined all genes differentially expressed in NBs of the brain compared to the VNC in both species **(Figure supplement 31)**. However, this list was extensive and when we tested some of the genes by FISH, we found many with a panneural pattern of expression such as *NK7.1*, *CG9932*, *mre11*, *Tob*, *tna*, or *tai* (**Figure 5)**. To make our analyses more sensitive, we focused only on TFs, used the robust PyDESeq2 tool (76), and removed the sensory NBs, midline NBs, mesodermal/muscle progenitors, PGCs, and embryonic hemocytes, from the datasets. Indeed, in tests we found that including even a small subset of non-NB cells in the analysis enhanced the detection of false positive genes (**Figure supplement 31**). This approach revealed 78 TFs in *Tribolium* and 137 in *Drosophila* to be differentially expressed in brain NBs (**Figure 4**). Among them, we found 27 brain-specific TFs shared by the two insects: *CG5516, D, Dll, Fer1, Hmx, Optix, Phf5a, Rx, TfAP-2, ac, ase, bi, croc, dac, disco, dmrt99B, erm, fd102C, ffih, hbn, oc, run, scro, tll, toy, tup, wor* (**Figure 4 and Figure supplement 29**). Confirming the validity of our analysis, many of these shared brain TFs were already known to be expressed in procephalic NBs such as *Optix, Rx, erm, fd102C, hbn, toy, oc or tll, among others.* Interestingly, we also identified TFs that had not been confirmed or associated with embryonic brain NBs before: *Hmx, TfAP-2, dmrt99B, CG32532*, *fd59A*, *CG15696,* and *Fer1*.

**Figure 5.**
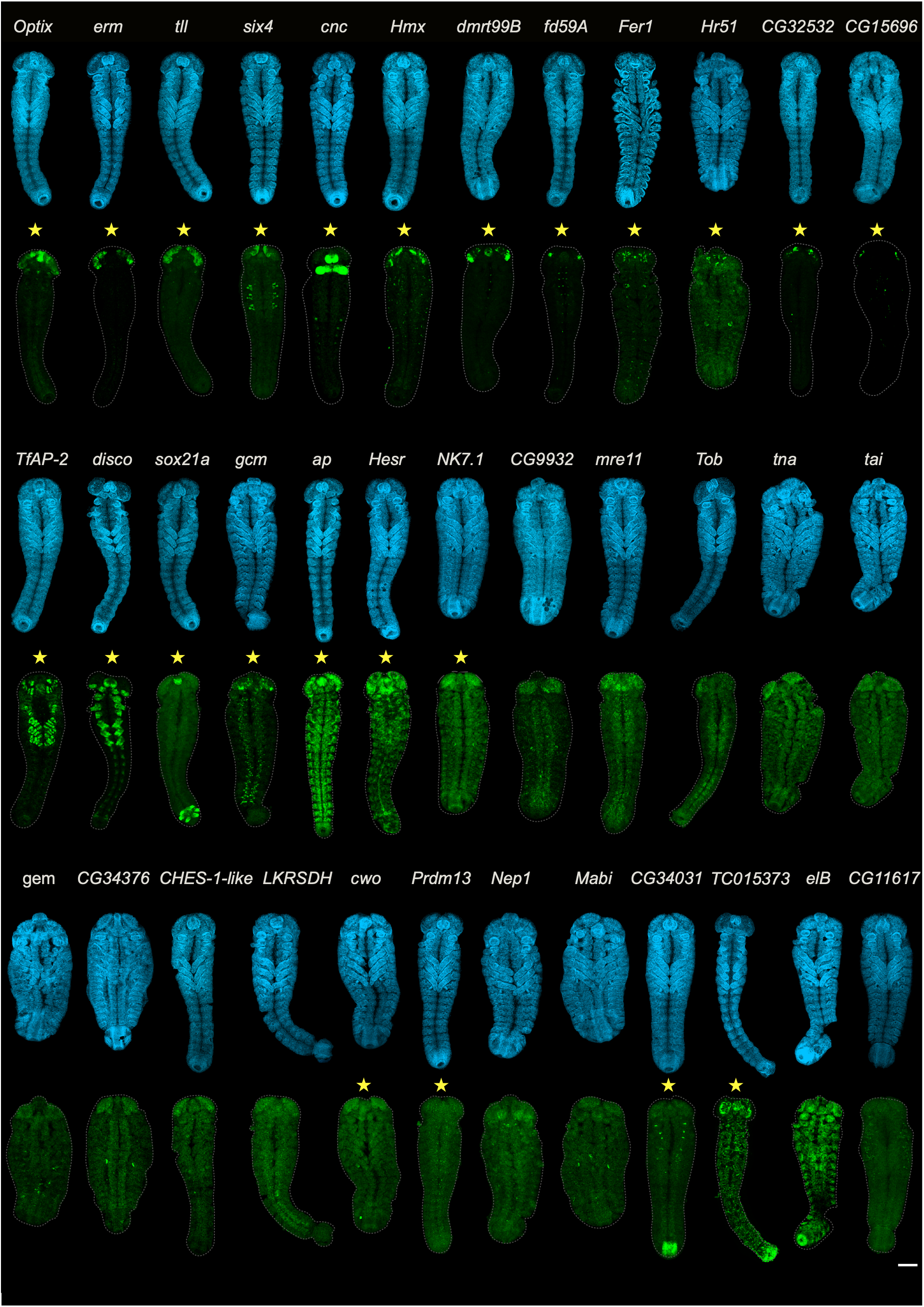
FISH to identify expression patterns of selected brain TFs in *Tribolium*. Overview of the expression of selected brain TFs by FISH (green) in elongated germband stages (see nuclear staining in blue). In **Figure supplements 37-92** more stages of these genes and the expression of several more genes are documented. Here, TFs are shown that were selected from the DE analyses of all genes, in addition to genes that were selected from the DE analysis of TFs. All depicted genes showed co- expression with the NB marker *asense*. Besides examples for very specific brain patterning genes (first row), we show genes with more complex patterns (second row). Besides well-known genes such as *Optix*, *erm*, *tll*, and *Six4* we also found genes that were not known to be present in brain NBs such as *Fer1*, *CG32532*, *CG15696*. In the third row, genes are shown that seem to be expressed in the entire CNS but were detected as brain-specific in our analysis. Some of them showed a higher expression in brain compared to VNC explaining their assignment in our analyses. We accepted this degree of false positive genes in our analysis in order to not lose any brain specific TF. Scale bar 100 μm.

To assess the reliability of the gene set identified by single cell sequencing and to gain information about the spatial expression of the novel brain TFs, we determined the expression of a subset of 24 TF candidates in *Tribolium*. Based on their expression in the BDGP *in situ* database (77) 8 TFs that in our analysis were detected only in *Drosophila* (*Six4*, *cnc*, *CG15696*, *Sox21a*, *NK7.1*, *cwo*, *CG34031, ap)*, 8 TFs found only in the *Tribolium* analysis (*fd59A*, *Hr51*, *CG32532*, *gcm*, *Hesr*, *Prdm13*, *TC05373*, *Pfdn2)*, and 8 TFs present in both insects (*Optix*, *tll*, *erm*, *Hmx*, *dmrt99B*, *TfAP-2*, *disco* and *Fer1)* (**yellow stars in Figure 5**).

Confirming our approach, many genes had brain-specific expression (*Optix*, *tll*, *CG15696*, *fd59A*, *CG32532*, *erm*, *Hmx*, *dmrt99B*, *Fer1*). Several genes were expressed in the brain and the appendages (*disco*, *TfAP-2, six4*, CG34031), but their posterior expression was not in NBs. Notably, some of these genes showed an early broad brain expression, which later split into discrete domains (e.g. *Optix, tll, six4* and *Hmx*) while others had very confined patterns from early stages onward (e.g. *fd59A, CG32532 and CG15696*), indicating different roles in brain patterning. Several of the tested genes were expressed in both the brain and the VNC (e.g. *ap*, *NK7.1*, *cwo, Hesr*), although we noted that some of them seemed to have a stronger expression in the brain. Taken together, the expression data indicates that our analysis successfully enriched for brain NB TFs but did not fully exclude genes expressed in the VNC.

In order not to miss brain TFs (i.e. to avoid false negatives), we regarded a certain level of false positives as acceptable. We also noted that the species-specificity was not confirmed, as all tested “fly-specific” factors were expressed in *Tribolium*, too (see below). In **Figure 5**, we present late-stage embryos for some of the genes that we tested by FISH. For all those genes, we confirmed co-expression with the NB marker *asense* in the brain of *Tribolium* for at least one stage (see **Figure supplement 37 to 85** for more stages and more genes that we tested for diverse reasons).

#### Conserved TFs in insect brain neuroblasts

From the brain-specific TFs, we selected seven genes for further analysis that had either not been connected to embryonic brain NB patterning or had not been well-studied before. In our analysis, four genes (*Hmx, TfAP-2, dmrt99B*, and *Fer1*) were detected in both species, *CG15696* was predicted only in *Drosophila*, and *CG32532* and *fd59A* only in *Tribolium* (**Figure 6A**). For these genes, we also determined the expression patterns in *Drosophila* and found that all had similar brain-specific expression as in *Tribolium* **(Figure 6B and Figure supplements 86-92)**. This shows that our approach of studying two insects increased the overall sensitivity. None of the genes that were found only in fly or beetle were really “species-specific” such that we assume that differences e.g. in dynamics of expression, in embryonic development, or technical issues influenced the sensitivity of detection. This also means the number of brain TFs conserved in both species will be much larger than the 27 TFs from our analysis (**Figure 6A**). *Hmx*, *CG15696*, and *CG32532* were expressed exclusively in embryonic procephalic domains, while *dmrt99B*, *fd59A*, and *Fer1* had additional patterns more posteriorly in advanced embryonic stages, and *TfAP-2* showed additional expression in the leg primordia. Lack of co-expression with *asense* indicates that these posterior expression domains are not in VNC neuroblasts. The gene *Hmx* is expressed rather broadly in the protocerebral neuroectoderm from early stages onwards and was found in several NBs (**Figure supplement 42**). The expression of the genes *dmrt99B, fd59A, CG15696 and CG32532*, in contrast, was restricted to very small procephalic regions from early stages onwards and were expressed only in a few NBs (**Figure supplements 43, 44, 47 and 48)**. *TfAP-2* was expressed in a few NBs in the head of the insects, but in addition showed non-NB expression in the legs (**Figure supplement 49**). This pattern combination was also found in *six4* and *disco* (**Figure 5**). *Fer1* started to be expressed late in embryonic development in only a few spots (**Figure supplement 45**). We comprehensively documented the patterns of expression in *Tribolium* and *Drosophila* throughout embryonic development. These images and higher-resolution images are found at https://doi.org/10.25625/QLDOOT.

**Figure 6.**
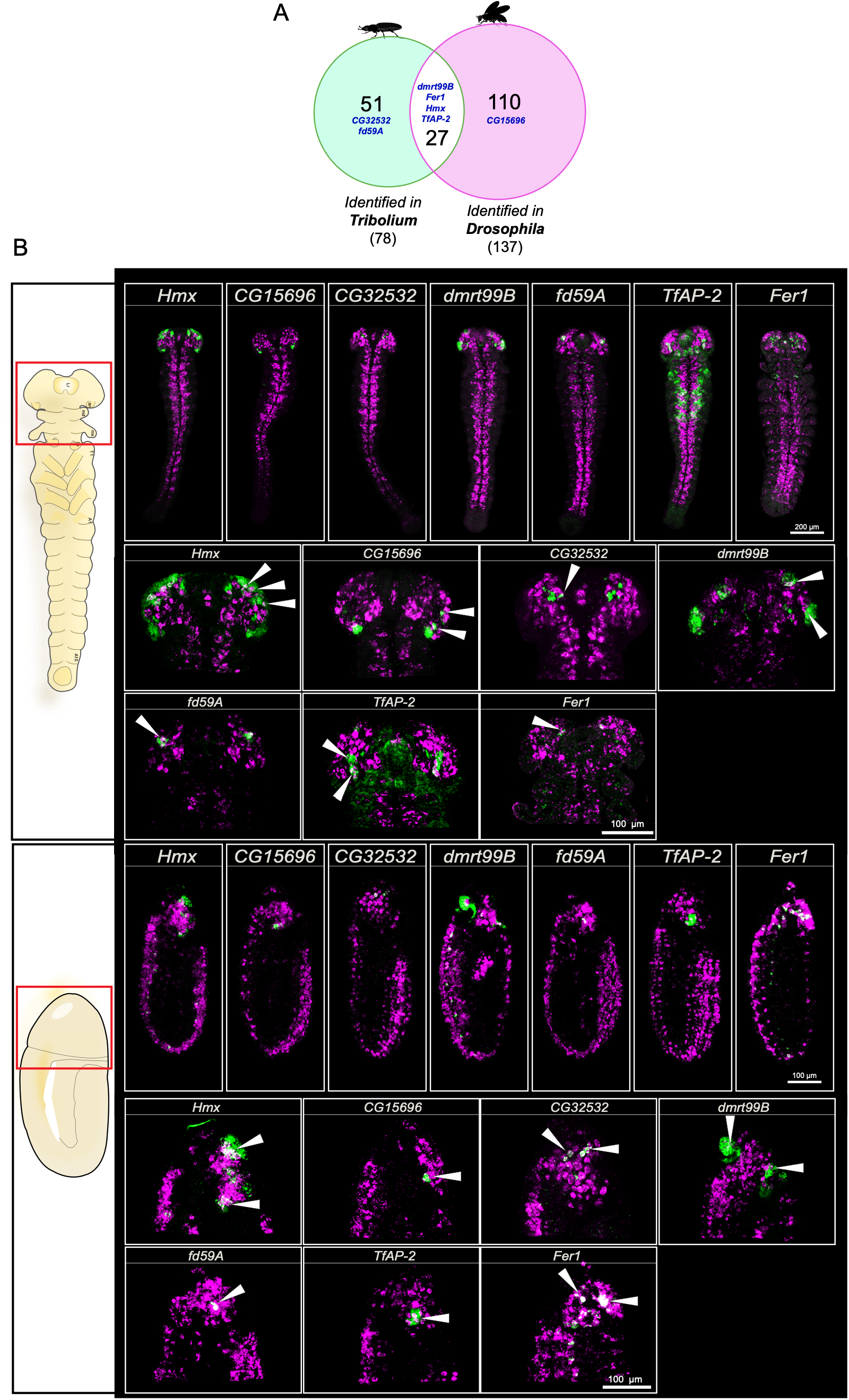
Selected TFs expressed in insect brain NBs. (**A**) Euler diagram of the transcription factors differentially expressed in brain NBs. 78 TFs were identified in *Tribolium* and 137 TFs in *Drosophila*, but 27 of them were identified to be the same in both insects. Based on our FISH analysis, also the TFs identified in only one species were expressed in brain NBs of both insects. Hence, we consider all 188 genes to represent the core of insect brain NB specification. Additional analyses are required to prove whether truly species-specific genes are among these genes. The gene names in blue represent the TFs that we tested and found to be expressed in brain NBs. (**B**) FISH staining for the NB marker *asense* (pink) and the TF candidates (green) in *Tribolium* and *Drosophila* embryos. The schematic embryos on the left show a ventral view of a stretched-out *Tribolium* embryo (top), while the *Drosophila* embryo is shown in lateral view. In both, the head anlagen containing the brain neuroectoderm is up and marked with a red square. All tested genes revealed similar expression in both species. The close-ups (corresponding to the red squares) are snapshots to display some of the brain NBs co-expressed with the TFs (white arrowheads). Since the co-expression in the neuroblasts since to be highly dynamic depending on the stage of development and the brain region, we encourage the reader to review the Z-stacks for each gene and embryonic stage provided in this work to spot further details. There, we also provide additional images that show other aspects of gene expression not in NBs (i.e. not co-expressed with *asense*) but in other tissues, mostly the epidermis.

#### Knock-down of brain-specific genes suggests brain neuroblast identities are given by TF cocktails rather than single master regulators

We wondered if these newly identified and highly conserved brain TFs had a broader function in brain patterning. We selected the earliest and most widely expressed gene (*Hmx*), one of the most specifically expressed genes (*CG15696*), and one gene with brain and leg expression (*TfAP-2*) for functional tests. First, we checked for a role in neuroectoderm patterning. As head cuticle and brain are derived from the same epithelium, we used the bristle pattern on the head cuticle as a sensitive readout for neuroectoderm patterning (33, 78, 79). However, RNAi knockdown of these genes in *Tribolium* caused no defects in either repetition (1 μg/μL and 2 μg/μL, respectively) with two different dsRNA preparations (**Figure supplement 93A and B**). The fact that most larvae had shortened legs in the *TfAP-2* knock-down in both repetitions indicated that we had no systematic problem with RNAi (**Figure supplement 93F, G, and H**). The identification of minor phenotypes in brain development is more challenging for several reasons. Given the complexity of the brain, a plethora of markers would have to be used to cover all aspects. Further, neural identity is specified by combinations of TFs such that the loss of one factor will lead to misspecification rather than the loss of lineages. Hence, the resulting changes are likely to be mild and may not affect the gross morphology of the brain, the cell bodies, or the macrocircuitry. In order to identify brain phenotypes, we used transgenic lines marking subsets of neurons that contribute to certain brain structures: The GEKU enhancer trap line G10011 marks neurons that express *shaking hands* (80) and the *foxQ2-5’-line* marks *FoxQ2*-expressing cells (34). These two lines mark the central complex and other brain structures. Our third reporter line marks the mushroom bodies (*MB-green* line) (54). We tested all lines with the genes *Hmx* and *CG15696* but found no phenotype when analysing the overall brain morphology (using synapsin staining) or the marked cell clusters and their main projections in first larval instar brains (**Figure supplement 94**). We conclude that these TFs are likely not individually required for the development of entire lineages and their macrocircuitry. Either they contribute to specification in dimensions that we had not screened for (such as changes in neurotransmitter content; modifications of microcircuitry) or their role is buffered by other TFs that are part of the cocktail that specifies NB identity.

## Discussion

We performed single cell sequencing of the embryonic neuroblasts of the two most advanced genetic insect models, which allowed us to carry out a cross-species single cell transcriptomic study to reveal the NBs conserved molecular and cellular landscape. This resulted in a dataset of more of nearly 70,000 NBs from *Drosophila* and *Tribolium*, that we classified into VNC and brain NBs (**Figure 1 and 2**). We identified the repertoire of transcription factors that are enriched in VNC and brain NBs (**Figures 3 and 4**) revealing that their transcriptional programs are distinct in both *Drosophila* and *Tribolium*. This suggests a profound divergence between the insect brain and posterior CNS reflecting recent reports in vertebrates (81). Finally, we identified novel TFs marking brain NBs and determined their spatial patterns of expression during the embryonic development (**Figures 5 and 6**). Our data contribute to understanding the ancestral developmental program and molecular adaptations that generate the diversity that we see in insect brains.

### Challenges of cross-species single-cell analyses

Cross-species comparisons are prone to biases introduced by differences in obtaining the samples, the sequencing methods, and the bioinformatics pipelines applied. Indeed, we found different numbers of brain TFs identified in *Drosophila* and *Tribolium* (**Figures 4 and 6**), which might be due to either biological differences or methodological bias. Bioinformatically, we used the identical bioinformatics pipeline for both datasets (**Figure supplements 5-18**) such that we expect this to reduce bias. However, we were not able to use the same method for the generation of the cell suspensions. In *Drosophila,* we fixed embryos and stained NB with the *wor*-antibody. We then produced nuclei suspensions that were sorted and finally sequenced NB nuclei (**Figure supplement 1**). In contrast, for *Tribolium*, we used a transgenic line with mostly cytoplasmic tGFP in NBs (55) and performed single-cell RNA sequencing after isolation of living NBs rather than nuclei (**Figure 1B**) (**Figure supplement 2**). Actually, we tested the fly *wor-*antibody, which unfortunately did not work in *Tribolium* and another antibody marking NBs is not available for *Tribolium*. We also tried isolating NB nuclei using an antibody targeting tGFP expressed in our transgenic line. However, after dissociation, the fluorescence of the nuclei was too low for sorting and enrichment. These differences in the generation of the cell suspension surely influenced our results and might limit the comparability across species. We think that this might be a reason that some genes were found only in flies and others only in beetles while all tested genes were actually expressed in both (**Figure 6**). We realized that the bioinformatics analyses of the *Drosophila* data led marginally sharper results. We speculate that the isolation of fixed and marked nuclei may be less prone to artefactual gene expression, which is known to occur in living cells under stress imposed by mechanical isolation and sorting. This consideration may become even more important when neurons are to be sequenced, as they would probably need to be treated more harshly to isolate them from their tight interconnections within the brain.

We found it to be of key importance to reduce the bias stemming from the different quality in omics resources and annotations between our model systems. *Drosophila* is clearly the model system with the best omics resources while *Tribolium* is among the most prominent runners-up with a great amount of omics data available (50, 82). Still, the available data was not sufficient for our comparisons. First, the available orthology inferences from OrthoDB had assigned only 5,000 genes to be orthologous between *Drosophila* and *Tribolium* (83). In contrast, our own prediction of orthologous genes focused specifically on *Drosophila* and *Tribolium* led to the identification of 9,000 orthologous genes (66). The enhanced number of well-determined orthologs surely increased the sensitivity of our comparison and our orthology list will be valuable for future comparative approaches between flies and beetles. Second, the more comprehensive knowledge in *Drosophila* may bias comparative data. For instance, we first performed our analysis based on beetle orthologs of known TFs from flies. However, the *Tribolium* TF-list was much shorter because beetle specific TFs were not covered by this approach. Indeed, in the analysis based on these TF lists, we detected worryingly more brain TFs in the fly than in the beetle. Therefore, we used an unbiased *de novo* prediction of TFs for both, *Drosophila* and *Tribolium* (74), where we found similar numbers of TFs in *Drosophila* (958) and *Tribolium* (868) (**Figure supplements 28 and 29A-C**). Indeed, the subsequent analyses led to a much more balanced number of brain TFs detected in either species. Still, the number of differentially expressed brain TFs is smaller in *Tribolium* (78) compared with *Drosophila* (137) and we do not know if this reflects a biological difference or whether the different quality of omics resources may still hamper the comprehensive detection in *Tribolium* (e.g. TFs missed in poorly assembled regions of the genomic sequence). In summary, for high-quality comparative single-cell studies, it is key to ensure that gene predictions and annotations are as comparable as possible. Depending on the depths of previous annotation efforts, it may even be required to completely re-annotate the genomic sequences (84).

### Evidence of profound divergence of the VNC and brain patterning

We find that spatial specification of anterior brain and VNC involves a strongly diverging TF set indicating different spatial patterning. This supports the notion of an ancient evolutionary separation of the anterior brain from the segmental VNC. Given that insect brains consist of three ganglia, we point out that our analysis comprises NBs from the anterior-most part, the pre-segmental region giving rise to the protocerebrum, including optic lobes, central complex, and mushroom bodies. In addition, the NBs from the Hox-free antennal segment forming the olfactory lobes are included, while the most posterior part of the brain, the tritocerebrum, was excluded due to Hox gene expression. While the tritocerebrum is regarded as a strongly reduced segmental structure, the antennal segment has, on one hand, similarities to trunk segments but also has a special position as the region behind the insect head boundary (i.e. the boundary corresponding to the vertebrate mid-hindbrain boundary) (4, 58).

The similarity of the genetic control of neuroectoderm, the NB specification and delamination and the principles of consecutive spatial and temporal patterning of NBs argue for the continuity of the entire CNS. In the extreme view, the brain would merely be an anterior specialization of the CNS with a higher number of NBs and a small number of brain-specific type II NB types. Reflecting this view, the TFs known from the VNC had been used to map the identities of brain NBs (12, 13, 22, 26–28). Conversely, a number of genes from the anterior gene regulatory network (aGRN) had been shown to be expressed in the anterior brain in most animals, lending support for the notion of a more ancient evolutionary divergence (4, 32, 33, 85, 86). Our unbiased identification of a vast number of brain-specific TFs lends strong support for the latter view, which had been based on the study of candidate genes (33, 87). Very recently, a study using vertebrate brain organoids also found a fundamental divergence of progenitors. Very early in gastrulation different progenitors exist in the fore/midbrain neuroectoderm compared to the hindbrain ectoderm, which diverge at the level of the chromatin landscape (81). Some have used the molecular divergence of the brain and the more posterior CNS to suggest that both parts have evolved separately and have later fused to form a chimeric brain(6). Our data strengthens the argument of profound divergence. However, given that several basic cellular and molecular mechanisms of neurogenesis are identical in both regions, we prefer the view that the CNS evolved in the last non-segmented common ancestor including a basic spatial separation. When the evolution of segmentation and somitogenesis diversified the posterior part of the animal, parts of the non-segmental CNS were multiplied to form the segmentally reiterated part of the CNS. The ancient anterior CNS patterning would then still be reflected in brain-specific TFs we see in extant animals. This model explains both, the similarities in the cell-biological mechanisms and the divergence of spatial specification.

### Unexpected divergence of NB Type II-like clusters

As expected, we found a cluster corresponding well to Type I NBs and this was also reflected by some of the most enriched genes (e.g. Notch pathway, *dpn,* and the apically localized *miranda*) (**Figure 3F, G**). However, the markers used for Type II NB identification led to three distinct clusters, where only one cluster fitted the expected pattern quite well (NBs II/INPs in **Figure 3**). Surprisingly, two additional clusters seemed atypical (NBs II/INP 1 and 2 in **Figure 3**; called atypical cluster 1 and 2 in this discussion). With respect to the canonical Type II NB cluster, we discovered several novel characteristics that merit attention: First, the NB Type I profile showed quite some similarities to the primordial germ cells (PGCs), while the NB Type II cluster was clearly distinct (**Figure 3F**). This indicates that Type I and Type II NBs may be more profoundly different than previously thought. Second, the Type II NBs showed high expression of many markers, which were also active in the other clusters (except PGCs). Therefore, Type II NBs were marked not so much by the expression of specific marker genes but rather by a higher level of expression of marker genes (**Figure 3F**). Third, we found *ypsilon schachtel* (*yps*) as a Type II NB marker, which is known for the posttranscriptional regulation of the *oskar* mRNA in oogenesis and maintenance of germline stem cells but had not been related to NB biology before. Our finding indicates that *oskar* could act in holometabolous NBs, which has not been described in flies so far but would be in line with data from the hemimetabolan *Gryllus bimaculatus* (88, 89). Given the absence of reports of *oskar* expression in fly NBs, it would be interesting to check this aspect in *Tribolium*, which has often proven to reflect the ancestral biology better than *Drosophila* (90–92). Fourth, in our results, the Toll/NF-κB and cGAS-STING pathways are most strongly correlated with the NB Type II cluster (**Figure 3G**). Especially intriguing is the latter pathway, which is an immune pathway that had not been linked to NB biology before. Overall, with respect to pathway involvement, the Type II NBs were more similar to sensory and midline NBs than to Type I or the other Type II clusters, which adds to the notion of unexpected divergence of NB types.

The atypical cluster 1 stands out with respect to the many pathways covered by its expression profile: It shows the second highest relation to the Notch pathway (second to Type I NBs) and is expressing components of almost all pathways except for the IMD pathway (**Figure 3G**). This is also reflected in the marker genes, which cover components of the EGFR, Wnt (*pan*), hedgehog (*ci, ihog*), Hippo, and PCP (*fat, dachsous*) pathways (**Figure 3F**). Strikingly, the planar cell polarity (PCP) pathway is recovered by two components (*fat* and *dachsous*), indicating a role of planar polarity (typically found in epithelia) in this clusters’ biology. This is unexpected as Type I NBs are known to develop largely cell-autonomously after delamination (8, 9). In contrast to that paradigm, the astonishing degree of pathways potentially active in atypical cluster 1 cells indicates that they are integrated into a cell-non-autonomous regulatory environment. Our data also relates the *ovo/shaven baby (svb)* gene to neural progenitor biology for the first time. *ovo* and *svb* are distinct isoforms of the same gene, but we did not distinguish which of them was expressed in the NBs. *ovo* has been well described with respect to oogenesis, and accordingly we found it enriched in PGCs. In line with the hypothesis of a novel role in fly brain NB biology, in a large in-situ-screen performed in flies, *ovo/svb* was found to be expressed in several distinct clusters in the late dorsal embryonic brain neuroectoderm (93). Further analyses are required to find out which isoform is expressed in NBs (94, 95).

The atypical cluster 2 is marked by comparably weak differential expression of the respective marker genes (**Figure 3F**). The enriched *fatty acid binding protein (fabp)* is active in glia and involved in neural development (96), while *pebbled (peb)* is a known tissue-specific transcription attenuator, which is active in eye development(97, 98).

What cell type or stage could these atypical clusters correspond to? Both atypical clusters show a strong link to the insulin pathway, which regulates growth and reactivation of quiescent NBs (99, 100). Hence, both likely represent growth or transition stages. The known differential activity of the Notch pathway (high in Type II NBs, downregulated in immature INPs and re-activated in mature INPs)(63) would argue for atypical cluster 1 representing mature INPs while atypical cluster 2 could represent immature INPs. The fact that the marker gene expression in cluster 2 is comparably mildly different from the other clusters supports the hypothesis of an immature stage that is transcriptionally not yet well-defined. If this assignment is correct, mature INPs (i.e. atypical cluster 1) would surprisingly be integrated into a complex non-cell autonomous regulation environment. Alternatively, cluster 1 could represent NBs of the optic lobes (101). In postembryonic optic lobe development, several signalling pathways are known to be involved in the progression of NB specification from the neuroectoderm and their patterning (102–104). This hypothesis would also explain the expression of PCP pathway components, which usually act in epithelia. This hypothesis could be a starting point to study development and biology of embryonic optic lobe NBs, which so far have not been well-studied.

With respect to other NB clusters, we note that the midline NBs showed the strongest connection to the IMD pathway. Hence, we identified two different immune pathways to be linked with different NB types: the cGAS-STING pathway with Type II and the IMD pathway with midline NBs. To our knowledge an involvement of immune pathways in NB biology has not been described before but IMD seems to be involved in damage induced regenerative proliferation of a variety of cells including glia (105).

### Novel TFs linked to brain neuroblast development

In contrast to the VNC, the spatial specification of brain NBs remains enigmatic and the regulatory code for the development of higher order brain centres remains scarcely understood. While an extensive co-expression map of TFs does exist (27, 28), this list was inspired mainly by knowledge from the VNC. Based on conservation across Bilateria, additional genes had been identified for insect head and brain patterning (4). Hence, both approaches relied on candidate genes rather than a comprehensive unbiased search for brain TFs. Our study was aimed to fill this gap by the first comprehensive sequencing of embryonic NBs in holometabolous insects. The 188 brain-specific genes shown in **Figure 6A** were identified in either one or both species but based on our control stainings, most or all of them will be expressed in both species (see above for possible technical reasons). This set represents the first unbiased genome-wide list of brain-TFs of the embryonic insect brain. Indeed, besides the expected genes, we found several brain-TFs that have not been well-studied before or have not been linked to brain development at all. Studying these genes will provide novel insights into insect brain development. Below we discuss some TFs, which we consider especially interesting to follow up.

First, we consider *Hmx* as a very interesting gene, because it is expressed from the earliest stages onwards, its later expression probably covers anlagen of higher order brain centres and is well-known for neural patterning in vertebrates. It is an ancestral homeobox TF reported to be conserved among insects, bilaterians, and cnidarians, with conserved regulatory sequences (106). This gene has multiple paralogs in vertebrates, while a unique gene is present in insects, and the fly *Hmx* is able to compensate for the roles of *Hmx1*, *Hmx2*, and *Hmx3* in a murine model (107). The expression of *Hmx* has been reported in lamprey, mouse, chicken, *frog,* zebrafish, Ciona, worms, and shark, where it is expressed in the CNS, head regions, eye, and the cranial, geniculate, and trigeminal ganglia (107–113). In insects, the expression of *Hmx* is reported for *Drosophila* embryonic development. It starts to be expressed at the blastoderm stage as to dorsolateral stripes at the anterior pole, and at stages nine to ten, several patches of expression appear. At this point, *Hmx* is co-expressed in some NBs. At stage 11, *Hmx* is also expressed along the VNC domains, but its function there is unclear (114) (**Figure 5, 6**, and **Figure supplement 42)**. Among other structures, these domains are close to or cover the insect placode (with evolutionary relationship to vertebrate placodes) and/or the location of Type II NBs, the eye and the mushroom body anlagen (4, 115). Second, *fd59A* seems interesting because it marks only one or very few brain NBs. It is a forkhead box (FOX) transcription factor found in all arthropods (116, 117). In flies, it is involved in apoptosis in spermatogenesis, egg-laying behaviour, neural maturation, and as a terminal neuronal selector in the larval optic lobe (118–121). Its embryonic expression has been described in *Drosophila*, *Tribolium, Glomeris, Parasteatoda, and Euperipatoides* (116, 119, 122). In all of them, *fd59A* appears at the most anterior domain of the head, and then the expression expands in this area, but is still localized at two spots in the head. At advanced embryonic stages, *fd59A* becomes also expressed along the VNC. We also detected the expression of *fd59A* in the stomodal region of *Tribolium*. In *Drosophila,* there is only one extensive study of *fd59A.* They describe embryonic and larval expression and find *fd59A* to be co-expressed with the *Nitric oxide synthase* (*Nos*) in Hb9 and octopaminergic neurons. Knockdown led to a reduction of laid eggs leading to the hypothesis of an involvement in mating behaviour. However, the number of neurons remained the same after *fd59A* knock-down (39, 119) (**Figure 5, 6**, and **Figure supplement 44)**. We find that in the brain neuroectoderm, its expression seems to be close to the mushroom body anlagen. The fact that NO plays crucial roles in many aspects of brain development and function, together with our finding that it marks only a few NBs, makes it an excellent candidate to ponder the origin of Nos expressing neurons. Third, *Fer1* is a basic helix loop helix (bHLH) transcription factor uncovered in *Drosophila* screening studies in endocrine cells and olfactory neurons, but *Fer1* has not been related to NBs so far (39, 123–127). The embryonic expression and function remain unknown in any insect. Our data suggests that *Fer1* is likely involved in early neuronal development since it is expressed in several patches in advanced embryonic stages (**Figure 5**, **Figure 6**, and **Figure supplement 45)**. Interestingly, we found co-expression with probably only one median NB at advanced stages (**Figure 6**), which could make it a very specific marker for one lineage or a small subset of neurons. Fourth, the gene CG32532 (The *Tribolium* ortholog is called *shaking hands, skh*) has recently been described with strong focus on *Tribolium* (80). It was described to be restricted to postmitotic neurons, which project into the central complex in both *Tribolium* and *Drosophila*. Based on its postmitotic and very specific expression, *skh* was interpreted to be a terminal selector of neuron identity. Our finding of *skh* expression in NBs at a similar location suggests that a part of the *central complex* neurons stem from this NB (**Figure 6** and **Figure supplement 48)**. Finally, we found the homeodomain transcription factor *CG15696* to be a brain-TF, which has not been studied before at all. We find it to be expressed in several very small spots of the brain neuroectoderm, one of which could be in the optic lobe or the antennal anlagen (**Figure 5,6** and **Figure supplement 47)**.

### A basis to understand the diversification of brain neuroblasts

In this work, we focused on detecting the conserved core of insect brain TFs and we provide novel entries into studying the development of the insect brain. Importantly, this dataset is also a basis for searching for the genetic bases of evolutionary brain diversification, which is known to involve at least differences in brain NB numbers among insects (13). One approach would be to extend the search for species-specific changes of the brain TF-set. On one hand, one should test the other genes from our list of 188 genes but one could also use our data to create a more extensive collection of candidate genes to be tested by FISH subsequently. A second approach would be to compare the expression of conserved brain TFs in more detail to reveal differences in expression timing or pattern. Finally, the data could be used to determine the cocktails of TFs in certain NBs to subsequently identify differences across species. Actually, our failure to see strong phenotypes upon knock-down of the three tested genes may indicate that interfering with one TF might lead to fine grained divergence rather than changes in gross morphology.

## Materials and Methods

### Animal husbandry

The *Tribolium* wildtype strain *San Bernardino* (SB) or the *vermilion^white^*, strain carrying a mutation in the *Tc-vermilion* gene were used (128). The beetles were reared at 32 °C with 40 percent humidity in whole-grain flour type 1050 and transferred to wheat flour type 450 for embryo collections. We used the wildtype *Drosophila* Oregon-R strain maintained at 18 °C on standard cornmeal agar. Embryos were collected on apple juice-agar plates coated with a thin layer of yeast paste at 25°C.

### Generation of the *Tribolium* GöGal41519 enhancer trap line

The line GöGal41519 was found in an enhancer trap screen and showed an embryonic pattern reminiscent of NBs while in the first larval instar a small number of longitudinal thoracic muscles were marked. The enhancer trap screen essentially followed the procedure of the GEKU screen (55). However, in contrast to the GEKU mutator, a *Tribolium* endogenous promoter was used (to increase expression) to drive Gal4delta (to increase usability beyond EGFP imaging) and an attP site was added (for further modification of the insertions). Moreover, *Tc-vermilion* (128) was used as eye marker to avoid the 3XP3 driven fluorescence in the glia, which had previously hampered analyses of brain enhancer traps. In the screen, a donor line carried the mutator with an enhancer trap that drove expression in the pupal abdomen (stock number and name: GB192; *Bauchbinde-donor-line*). The UAS responder drove turbo GFP (tGFP) (stock number and name: GB104; *UAS-hsp-tGFP-6*) (51). The *Bauchbinde-donor-line* was first mated with the X-linked helper line M26 (129) (P1 generation; see **Figure supplement 2A** for mating scheme). A male of the P2 generation was mated with the responder line and in the F1, males were selected that still had black eyes (i.e. mutator still present) but had lost the Bauchbinde pattern (i.e. mutator at new location). Such males were mated with the responder line again. Animals carrying both the mutator (black eyes) and the responder (red eyes) were used to establish a stock, which was scored for new enhancer trap patterns at embryonic, larval, and pupal stages (see **Figure supplement 2B** for embryonic endogenous fluorescence and **Figure supplement 3** to observe the embryonic pattern of expression). The sequence of the mutator is available in GenBank accession number PZ173269. The insertion site was determined by iPCR as described previously (55) and revealed a position close to the *Tc-cyclin E* gene.

### *Tribolium* cell dissociation and cell sorting

We collected 35-50 hours old embryos and transferred them into a washing basket. We washed the embryos by submerging the basket several times in Milli-Q water to remove traces of flour, and subsequently dechorionated the embryos with 5% bleach (DanKlorix Hygiene Cleaner) for 3 minutes under constant shaking. To remove all traces of bleach, we washed the embryos extensively with Milli-Q water. Henceforth, we kept all solutions, tubes, and embryos on ice to minimize cell stress and death. Using a paintbrush (Da Vinci watercolor brush, Art.Nr. 1526Y-0), we transferred the embryos into a 2 mL dounce homogenizer (Merck, D8938) prefilled with 1 mL of Schneider’s medium mixed with 1% BSA (Thermo Fisher Scientific 21720024 and 15260037). We then mechanically dissociated the embryos by using approximately 15 strokes of the pestle A. We decanted the homogenate into a 1.5 mL vial and centrifuged it at 50g for 3 minutes at 4 °C to remove chunks of tissue. We decanted the supernatant into a new 1.5 mL vial and centrifuged it at 300g for three minutes at 4 °C. We discarded the supernatant and resuspended the pellet in one millilitre of Schneider’s medium mixed with 1% BSA. We filtered the suspension with a 20 μm cell strainer (PluriSelect 43-10020-60) coated with a thin layer of 1% BSA and centrifuged again for three minutes at 4°C. Finally, the cell pellet was resuspended in 50 μL of cold Schneider’s medium mixed with 1% BSA supplemented with 0.8% protector RNase-inhibitor (Roche 3335402001) and 1x Protease Inhibitor Cocktail (Promega G6521). The vials containing the cell suspensions were placed on ice and immediately transported to the FACS facilities. Using the cell sorter SH800S (Sony Biotechnology Inc) with a 100-μm microfluidic sorting chip (Sony LE-C3210), cells were sorted for one hour. To set the gating parameters for FACS, we used unstained cell suspensions from wild-type *Tribolium* embryos as a negative control and DAPI stained wild-type embryos (700 ng/mL for 15 minutes) to set the gating for live versus dead cells. Once the gating parameters were set, we started by analysing all events in our samples stained with DAPI using the forward scatter (FSC-A) and the side scatter (BSC-A) without gating. We then gated using forward scatter (FSC-A) and side scatter (FSC-W) to discard doublets. We then created a third plot using forward scatter (FSC-A) and fluorescent (DAPI), selecting DAPI-negative cells. Finally, these selected events were plotted again using phycoerythrin (PE) versus fluorescent (GFP) to exclude auto-fluorescent cells and to identify a small cell population of highly GFP-fluorescent cells **(Figure supplement 4B and 4C)**. We sorted the highly GFP-fluorescent cells into a centrifuge tube coated and prefilled with 50 μL of ice-cold Schneider’s medium mixed with 1% BSA.

### *Drosophila* nuclei dissociation and cell sorting

We collected embryos from the developmental stages 10 and 11, dechorionated them using 5% bleach for one minute, and thoroughly washed them with water before transferring them into a microcentrifuge tube. Embryos were then fixed with a mixture of equal parts of heptane and Dithiobis(succinimidyl propionate) (DSP) (1mg/mL) for an hour in a rotary shaker (130). Followed by three washes with absolute methanol, they were stored in methanol at −20 °C. Then, we performed antibody staining with RNAse-free reagents. We rehydrated the embryos with PBT in increasing concentrations mixed with methanol (25%, 50%, 75%) and washed them four times with PBT. Each wash lasted 10 min. Embryos were incubated twice for 15 minutes in blocking solution (PBT + BSA 1%). The primary antibody Rat-anti-Worniu (Abcam ab196362, Rat monoclonal IgG2a) was diluted at 1:250-1:500 in blocking solution and incubated overnight at 4 °C. Then, the sample was washed six times with PBT, 10 minutes each. Embryos were placed again in blocking solution twice for 15 min. The secondary antibody (Goat-anti-Rat-Alexa Fluor® 555, Abcam ab150158) was diluted in blocking solution and used at 1:500. The sample was incubated for four hours at 4 °C. Excess secondary antibody was removed by washing in PBT six times for 10 min. Epifluorescence microscopy confirmed staining **(Figure supplement 1A)**. For nuclei dissociation, the embryos were transferred into a dounce homogenizer (Merck, D8938) and washed three times with ice-cold PBS. PBS was discarded and 1 mL of dissociation buffer (PBS + BSA 0.04%) was added. The embryos were disrupted with the loose pestle (Pestle A). The suspension was transferred into a 1.5 mL vial. The douncer was washed with an extra 500 μL of dissociation buffer that was collected in the same tube. The sample was centrifuged for 3 minutes at 3000g at 4 °C, and the supernatant was discarded. The sample was resuspended with 1 mL of dissociation buffer and centrifuged for 3 min at 40g at 4 °C. The supernatant was collected in a new vial and centrifuged once more for 3 mins at 1000g at 4 °C. The supernatant was discarded and the pellet was resuspended in 1 mL of dissociation buffer. The suspension was sucked up and down 20 times with a 5mL syringe with a 22G x 2” needle. The cell suspension was filtered with a 20 μm cell strainer (Greiner Cat# 542120) and centrifuged for 3 minutes at 1000g at 4 °C. The pellet was dissolved in 400 μL of dissociation buffer. For cell sorting in a BD FACSAriaIII™ cell sorter, the samples were stained with DAPI and filtered with a 35 μm cell strainer. We first separated nuclei from cells using the forward scatter area versus the side scatter area (FSC-A vs SSC-A). Secondly, we used the forward scatter width versus the forward scatter area (FSC-W vs. FSC-A) to remove doublets. Finally, we plotted Alexa 555 versus DAPI and selected the neuroblast nuclei as the intersection of highly fluorescent events **(Figure supplement 1B)**. The selected nuclei were collected in 80% methanol. To reverse the crosslinks of the DSP fixation, nuclei were centrifuged for 3 min at 1000g, and the supernatant was discarded. Then, nuclei were resuspended in dissociation buffer with dithiothreitol (50mM) and incubated for 30 min at 37 °C. The sample was centrifuged for 3 min at 1000g, and nuclei were resuspended in dissociation buffer. The last centrifugation was repeated, and the final pellet was resuspended in 400 μL of dissociation buffer. Cells were finally counted in a Neubauer chamber **(Figure supplement 1C)** and diluted.

### Library preparation and single-cell RNA sequencing of *Tribolium* samples

We generated single-cell RNA libraries using the Chromium Next GEM Single Cell 3ʹ Kit v3.1 and a 10x Chromium controller (10x Genomics) following the manufacturer’s protocol instructions. The resulting cDNA libraries were quantified using a Bioanalyzer and sequenced in an Illumina NextSeq 2000 Sequencing Systems (Illumina, Inc.) to an average depth of 50,000 reads per cell.

### Library preparation and single-cell RNA sequencing of *Drosophila* samples

Single-cell RNA seq libraries were produced using Chromium Next GEM Single Cell 3ʹ Kit v2 and a 10x Chromium controller (10x Genomics). The cDNA libraries were quantified using a Bioanalyzer and sequenced in an Illumina HiSeq 4000 Systems (Illumina, Inc.) to an average depth of 30,000 reads per cell.

### Sequence alignment of *Tribolium* single-cell data

FASTQ files were analysed following the standard CellRanger pipelines (7.1.0). First, we used “cellranger mkref” to build a custom reference genome giving as input the *Tribolium castaneum* genome (Tcas5.2) and the transcriptome (OGS3) that included the tGFP sequence. To perform sequence alignment and produce the count matrices, we used “cellranger count” with the FASTQ files and the custom reference genome. The resulting matrices were aggregated with “cellranger aggr”.

### Sequence alignment of *Drosophila* single-cell data

FASTQ files were analysed using the PiGx pipeline (131). It aligned sequences using STAR, performed reads quality control, determined cell number, and produced the output matrices.

### Single-cell data quality controls, dimensionality reduction, and clustering

The matrices of *Tribolium* and *Drosophila* were analysed following the same pipeline for the sake of comparability. Data analyses were performed using the toolkit Scanpy (60). We loaded the matrices into an anndata object and identified the number of UMIs per cell and the percentage of mitochondrial and ribosomal genes. We removed cells that contained >10 percent of mitochondrial genes, >10 percent of ribosomal genes, <200 and >4,000 genes, and >50,000 UMIs. We also removed genes that were present in less than 10 cells (**Figure supplement 5-8**). We classified the cells as VNC neuroblasts if they contained gene counts for at least one Hox gene, and as brain NBs if they did not have gene counts for any of the Hox genes. We continued with data normalization and log transformation in Scanpy. Using further functions of Scanpy, we regressed out total counts, percentage of mitochondrial genes, and percentage of ribosomal genes. We then identified the 2,000 most informative genes and ran a principal component analysis (PCA). We generated a scree plot to rank the PCs and identified the most significant PCs. Based on this, we considered 20 PCs to calculate the number of neighbours and generated an unsupervised clustering using the Leiden graph-clustering method. The results are presented in UMAP and t-SNE plots at several resolutions (0.5, 1, 1.5, 2, 3, and 4) (**Figure supplement 9-14**). Finally, we calculated the top 30 marker genes for each cluster at resolution 1 (**Figure supplement 15 and 16**).

### Cell annotation of brain NBs

Unsupervised clustering of all NBs alone did not show a clear classification into subtypes (**Figure supplement 17 and 18**). Therefore, we subsampled the cells to retain only brain NBs and performed dimensionality reduction and unsupervised clustering. Then, to avoid mix signals and to make sense of the current knowledge of NBs, we manually checked in the *Drosophila* dataset the classical gene markers of NBs Type I, GMCs, NB Type II, INPs and labelled them accordingly (15, 132). When we observed unexpected clusters, we reviewed the top 20 gene markers to determine the cell type. In this way, we found clusters containing distinctive cell types of midline NBs, sensory NBs, or a mix of cells such as muscle and mesodermal progenitors, and embryonic hemocytes (**Figure supplement 19 and 27**). We attempted to annotate *Tribolium* NBs in the same manner. However, some of the classical gene markers were not recovered during the orthology prediction and with the markers available, we could not see a clear classification of the NBs as in *Drosophila*. In order, to achieve the annotation for *Tribolium* brain NBs, we concatenated the subsampled brain NBs datasets of *Drosophila* and *Tribolium,* performed data integration with Harmony, and used label transferring with scvi-tools and Scanpy (59–61).

### Pathway activity analysis in brain neuroblasts and other cells

To gain an overview of the pathway’s activity in the brain NBs and other cell types that we annotated, we downloaded the predefined set of core genes for signalling pathways from Flybase (72). Each set of genes contained several core genes that were grouped by pathway and duplicated genes within each pathway were removed. We then retained only the genes for each pathway that were expressed in our dataset. Pathway activity scores were calculated for each cell type using the Scanpy function *Score genes*. To assess pathway activity, we averaged the pathway scores by cell types, and the resulting mean pathway activity was visualized in a hierarchical heatmap using a row-wise standard scaling to normalize pathway activity.

### Gene ontology (GO) enrichment analysis in brain neuroblasts and other cells

We used the differential expression analysis performed in Scanpy using the Wilcoxon rank-sum test to identify up to 500 gene markers with adjusted p-values < 0.05 for our annotated cell types. These genes were then used to perform gene ontology (GO) enrichment analysis using g:profiler against the *Drosophila melanogaster* annotation database (133). We performed GO for two categories: biological processes and molecular functions. To make the visualization of this analysis simpler, we grouped the GO terms into broader GO slim-like functional categories. For the biological processes, we classified them into eight major categories: neuronal process, development, enzyme activity, immune response, transport activity, cell cycle, metabolism, and other. For the molecular functions, we used five major categories: neuronal process, cell cycle, enzyme activity, binding activity, and other. We used the enrichment p-value within each GO slim category to construct matrices. We then transformed p-values using −log10 scaling and normalized to a 0-1 range across all categories and cell types. We visualized the results in a hierarchical heatmap.

### *Tribolium* ortholog prediction

We predicted orthologous genes of *Tribolium* and *Drosophila* using OrthoFinder, the eggNOG 6.0 database, and in some cases phylogenetic trees, to address the equivocal orthology of important genes. The lists of orthologous genes are available in iBeetle-Base (66, 67).

### Prediction of gene transcription factors

For *Drosophila*, we combined available lists of transcription factors from the literature (**Figure supplement 28**) (15, 41, 70–73). However, there was no collection of transcription factors for *Tribolium*. To keep the analysis comparable across species, we did a *de novo* prediction using the deep learning tool DeepTFactor on the proteomes of *Drosophila* (dmel_r6.59) and *Tribolium* (Tcas5.2) (**Figure supplement 29**) (74). The predicted transcription factors for each species are available in our GitHub repository.

### Single-cell differential expression analysis

We used the Python libraries Scanpy, NumPy, pandas, seaborn, Matplotlib, SciPy, random, and PyDESeq2 (60, 76). Data analyses were performed independently for *Tribolium* and *Drosophila*. First, we loaded the h5ad matrix of the corresponding organism and removed mitochondrial, ribosomal and for the case of the brain NBs, we removed the cells that were annotated as midline NBs, sensory NBs, mesodermal and muscle progenitors, embryonic hemocytes, and primordial germ cells. We proceeded to identify all differentially expressed genes (DE) and differentially expressed transcription factors of the brain versus VNC neuroblasts by using pseudobulk samples with 5, 10, and 15 pseudo-replicates. (**Figure supplement 30 to 36**). We only considered the genes differentially expressed when the genes had a mean normalized count of at least 10 in order to remove lowly expressed genes with limited statistical reliability, an adjusted p-value threshold of 0.05, and absolute log2 fold change greater than 1.5. We standardized gene expression values across genes using a row-wise z-score normalization and plotted the gene expression of significant differentially expressed genes in a hierarchical heatmap.

### Embryonic fluorescent *in situ* hybridization (FISH)

RNA probes of around 800 bp were amplified by PCR from cDNA and cloned into the pJet1.2 plasmid. We used these vectors to synthesize RNA probes with Dioxygenin-UTP (Roche 11277073910) or Fluorescein-UTP (Roche 11685619910) haptens using the T7 RNA polymerase (Roche RPOLT7-RO) following the instructions of the manufacturer. Embryos were dechorionated with 50% bleach, fixed, and stored in absolute methanol at −20 °C. Prior to *in situ* hybridization, embryos were rehydrated in a dilution series of methanol in PBT (PBS + 0.1% Triton X-100) in decreasing concentrations (75%, 50%, 25%), followed by three washes with PBT. We proceeded with the *in situ* hybridization using Alexa Fluor™ 555, Alexa Fluor™ 647 (Thermo Fisher Scientific Inc. B40955 and B40958) and DAPI. After the final wash, we removed the washing buffer, added 50 μL of Vectashield Antifade Mounting Medium (Vector Laboratories H-1000-10), and stored the stained embryos at −20 °C. The detailed protocol has been previously described (134).

### RNAi

We cloned *Tribolium* orthologs of *Hmx* (*TC012136*), *CG15696* (*TC015928*), and *TfAP-2* (*TC009922*) into the Pjet1.2 vector. We used primers with T7 overhangs to produce PCR amplicons for dsRNA production using the 5X Megascript T7 kit (Ambion). We performed parental RNAi using 1-2 µg/µL of dsRNA as previously described (135).

### Larval (L1) cuticle analysis

After parental RNAi in the wildtype *Tribolium* strain and crossing, we collected the L1 larvae and performed clearing by embedding them in Hoyer’s medium mixed with lactic acid (1:1) in microscope slides that were then incubated at 55 °C overnight or until larvae was fully cleared (136). Larvae were covered with a coverslip and a minimum of 25 were examined under the microscope for overall phenotypes and head cuticle phenotypes.

### Larval (L1) brain antibody staining

We used *Tribolium* transgenic lines that mark brain structures of the larval brain: GB100-11 (marks the central complex partially), GB120 (marks the outline of the central complex), and GB176 (marks mushroom bodies). After parental RNAi in these lines and crossing, we collected embryos and let them develop until the L1 stage. We then dissected the brains and fixed them for 1 hour in 4% formaldehyde (Pierce™ 16% Formaldehyde (w/v), Methanol-free, Thermo Scientific™: 28908) and performed immunohistochemistry as previously described (137). Using the primary antibodies rabbit anti-GFP (Dilution 1:1000) (Invitrogen: A-11122) and mouse anti-Synapsin (Dilution 1:25) (DSHB: 3C11 (anti SYNORF1)). We used the secondary antibodies Alexa Fluor 488 and Alexa Fluor 555 (Invitrogen™: A-11070, A-21425). To stain nuclei, we used DAPI (Dilution 1:1000) (Invitrogen™: D1306). We repeated the experiments in at least three brains for each experiment.

### Sample mounting and microscopy

On a microscope slide, we placed the FISH-stained *Tribolium* embryos stored in Vectashield and proceeded to remove grains of yolk from embryos using a brush made with a pipette white tip and an eyelash. Afterwards, embryos were transferred onto a new microscope slide previously prepared with an adhesive paper circle and 1 μL of Vectashield, and if necessary, the embryos were arranged in a straight, flattened position. We covered the embryos with a coverslip of 1.5 thickness and gently pressed it using one eyelash brush. *Drosophila* embryos were mounted in the same way, but without flattening and removal of yolk. Confocal stacks were gathered using a Zeiss LSM 980 microscope, cuticles of larvae were acquired using a Zeiss AxioPlan 2 microscope using the software ZEN Blue 3.2.

### Image analysis and figure preparation

Image visualization and exporting were performed with the image analysis version of the software ZEN Blue 3.2 and Fiji (2.9.0) (138). Final figures were assembled using Adobe Illustrator and Adobe Photoshop (26.2.0).

## Declaration of use of AI and generative AI software

During the preparation of this work, the authors used Chat AI, a webservice provided by GWDG, the IT provider of the Max Planck Society (MPG) and the University of Göttingen, to troubleshoot some of the code. The authors reviewed the contents and outputs and take full responsibility for the content of this manuscript.

## Acknowledgements and funding resources

NC is grateful for the support provided by the International Max Planck Research School for Genome Science (IMPRS-GS). With respect to the Gal4 enhancer trap screen, we thank Johannes Schinko for providing clones and lines and Elke Küster for performing the screen and doing initial screening. We also thank Lena Reim, Artim Lange, and Claudia Hinners for their support in producing some of the stainings.

## Contributions

GB, NC, AV, and RZ conceived the study. AV and NC performed sequencing experiments in flies and beetles, respectively. NC performed computational analyses and all subsequent experiments. GB and RZ supervised the experiments. GB and NC wrote the original draft of the manuscript. RZ and AV revised and commented on the manuscript.

## Data sharing

The raw sequencing files and raw count matrices are available in the NCBI database Gene Expression Omnibus under accession number:<u>GSE337265</u>. Processed matrices are available at: https://doi.org/10.6084/m9.figshare.33338799. Raw confocal images of the double whole mount *in situs* of the embryos of *Drosophila* and *Tribolium* produced for this study are archived in the following repository: https://doi.org/10.25625/QLDOOT

The Jupyter notebooks containing the code for the analyses presented in this work are available on GitHub at: https://github.com/SalmonellaIIB/Single_cell_neuroblasts.git.

## Preprint servers

https://www.biorxiv.org/content/10.64898/2026.08.27.747565v1

The copyright holder for this preprint is the author/funder, who has granted bioRxiv a license to display the preprint in perpetuity. All rights reserved. No reuse allowed without permission.

